# Flooding patterns shape microbial community in mangrove sediments

**DOI:** 10.1101/2024.07.24.604998

**Authors:** Mirna Vázquez-Rosas-Landa, Rosela Pérez-Ceballos, Arturo Zaldívar-Jiménez, Stephanie E Hereira-Pacheco, Leonardo D. Pérez-González, Alejandra Prieto‑Davó, Omar Celis-Hernández, Julio C. Canales-Delgadillo

## Abstract

**Background:** Mangrove ecosystems located in the tropics and subtropics, are crucial for regulating global weather patterns and sequestering carbon. However, they face threats from human activities like altered water flow and deforestation. While the symbiotic relationship between mangrove trees and surrounding microbes are essential for their survival, the impact of human activity on these microbial communities remains incompletely understood. We investigated how microbial communities change in degraded mangrove ecosystems due to loss of hydrologic connectivity, aiming to elucidate consequences and inform restoration strategies.

**Methods:** Employing 16S rRNA sequencing, we analyzed samples of sediment cores from conserved, moderately degraded, and degraded mangrove sites across dry and flood seasons at three sediment depths.

**Results:** Our analysis identified 11,469 Amplicon Single Variant (ASVs), revealing diversity loss correlated with degradation levels. Notably, we observed shifts in microbial diversity within sediment layers, with conserved sites dominated by Vibrionaceae in upper layers, potentially indicating urban contamination. In moderate-degradation sites, seasonal patterns emerged, with Halomonas and Marinomonas dominating the dry season and Exiguobacterium thriving during flooding. Interestingly, a community mainly composed of Firmicutes persisted across all degradation scenarios in deeper sediment layers, suggesting potential for ecosystem restoration. Our findings provide insights into microbial responses to human-induced stressors and highlight the role of core microbial communities in guiding restoration efforts for degraded mangrove ecosystems.

## Introduction

Mangrove ecosystems, found in tropical and subtropical regions worldwide (Lee et al., 2014), are vital components of coastal landscapes and play essential roles in global weather patterns, including carbon sequestration (Chatting et al., 2022; Song et al., 2023). Despite their ecological significance, these ecosystems face severe threats from human activities such as altered river flows and extensive deforestation (Akram et al., 2023). Within these ecosystems, microorganisms play key roles in sustaining ecosystem health and functioning, however, their role, particularly as a potential link for restoration, remains to be fully understood.

In mangrove ecosystems, microorganisms participate in nutrient acquisition, disease resistance, nutrient recycling, and stress tolerance (Palit et al., 2022; Holguin, Vazquez & Bashan, 2001; Akram et al., 2023; Bashan & Holguin, 2002; Vovides et al., 2011; McKee & Faulkner, 2000; Woodroffe et al., 2016). Mangrove trees and the microbial community interact mutually to support each other’s survival (Akram et al., 2023). For instance, mangroves shelter and nourish various microorganisms in their rhizosphere and root tissues (Purahong et al., 2019), and in return, these microorganisms facilitate nutrient cycling, such as nitrogen-fixation (Sjöling et al., 2005; Vovides et al., 2011), an essential nutrient often limited in coastal environments, thereby promoting mangrove growth and productivity. These diverse microbial communities also serve as buffers enhancing resilience to environmental stressors such as salinity, flooding, and disease (Lai et al., 2022). For instance, adding *Azospirillum* in restoration efforts promotes nutrient absorption and stress tolerance in mangrove roots by facilitating the colonization of diverse microbes (Domínguez-Núñez & Berrocal-Lobo, 2021). Additionally, a diverse array of bacterial genera, including *Agrobacterium, Alcaligenes, Arthrobacter, Bacillus, Enterobacter, Erwinia, Pseudomonas, Rhizobium, Serratia, Stenotrophomonas, Streptomyces*, and *Xanthomonas*, have shown protective effects against fungal and bacterial pathogens in various plant systems by colonizing infection sites, competitively excluding pathogens, and secreting antimicrobials (Bonaterra et al., 2022). Although reports of this phenomenon in mangroves are limited, similar mechanisms could operate within mangrove ecosystems. Leaning to the idea where mangrove trees and the surrounding microbial community form a resilient entity known as holobiont (Allard et al., 2020), challenging traditional views of organisms and emphasizing the interconnectedness between hosts and their associated microbial communities (Morris, 2018).

Recognizing the holobiont concept in mangrove ecosystems is key for conservation and management efforts. Conservation strategies should prioritize preserving mangrove species and safeguarding the diversity and integrity of associated microbial communities (Lai et al., 2022). For instance, it has been shown that different tree species, especially *Aviceina germinians* had different microbial communities associated (Gomes et al., 2014), and also core microbes and specialized microbiota to each tree has been observed (Wainwright et al., 2023), therefore, the success of the restoration is related to the microbial communities present in the roots of plants (Gomes et al., 2010). Integrating hydrological dynamics into restoration plans is vital, as water availability, sediment deposition, and nutrient cycling profoundly influence mangrove survival and regeneration (Pérez-Ceballos et al., 2020; Ellison, Felson & Friess, 2020; Akram et al., 2023). Thus, holistic restoration initiatives that consider the entire holobiont and incorporate hydrological factors are essential for ensuring the long-term resilience and sustainability of mangrove ecosystems.

Tidal fluctuations in mangrove ecosystems dictate crucial factors such as nutrients, oxygen levels, dispersal mechanisms, and salinity. These elements are essential for the well-being and distribution of mangrove species and their associated microbial community. Human-induced alterations to hydrological patterns can disrupt these ecosystems, leading to reduced tidal flushing, elevated salinity, sedimentation, and nutrient imbalances (Kamali & Hashim, 2011; Pérez-Ceballos et al., 2020). Conversely, prolonged flooding can raise water sulfide levels, harming mangrove trees (Pérez-Ceballos et al., 2022).

Hydrological dynamics also shape the distribution and composition of microbial communities in mangrove habitats (Mai et al., 2021; Thomson et al., 2022). Tidal inundation introduces diverse microbial populations from neighboring areas, enriching soil microbiota. Changes in hydrological patterns can disrupt these microbial communities, affecting nutrient availability, decomposition rates, and ultimately mangrove health (Pérez-Ceballos et al., 2020; Akram et al., 2023). Therefore, mangrove rehabilitation efforts should prioritize actions like restoring tidal connectivity and managing sedimentation to foster optimal conditions for mangrove growth and the establishment of healthy microbial communities.

Building upon the importance of microorganisms and hydrological dynamics in mangrove ecosystems, our study aims to investigate the impact of ecosystem degradation on microbial communities within mangrove sediments. Samples were collected in the Laguna de Términos, south Gulf of Mexico, from non-degraded mangrove, moderately degraded, and further degraded mangrove across two seasons and three sediment depths, with microbial diversity assessed using 16S rRNA sequencing. We hypothesize that increasing degradation will reduce microbial diversity, anticipating seasonal and depth fluctuations in composition. Through identification of key microbial species, our findings offer valuable insights to guide restoration initiatives amid anthropogenic pressures. Estero Pargo within Laguna de Términos, our study site, serves as an urban mangrove model to study these impacts, exhibiting notable changes such as fish farming expansion and tree mortality. This mangrove system’s zonation comprises red (*Rhizophora mangle*), white (*Laguncularia racemosa*), and black (*Avicennia germinans*) mangrove species. Estero Pargo provides a pertinent context for understanding the effects of human activities on mangrove ecosystems.

## Materials & Methods

### Study area

Laguna de Términos, designated as a Ramsar site of international and high biological importance, is currently experiencing several pressures due to growing industry and urbanization. The study area encompasses a mangrove forest situated within an estuary known as Estero Pargo on the southeast side of Isla del Carmen in Campeche, Mexico (18° 39’ 05” N; 91° 45’ 31” W and 18° 39’ 03” N; 91° 45’ 26” W, Supplementary figure 1). Estero Pargo, a tidal channel spanning approximately 6 km in length and averaging 14 m in width, covers a surface area of approximately 52 ha. The study area is located approximately 1-2 km away from the nearest urban settlements, with at least two untreated wastewater discharges entering Estero Pargo. Additionally, over the past three years, fish farming activities have expanded within the estuary, along with an increase in the size of a patch of dead trees. Consequently, Estero Pargo represents an urban mangrove that can serve as a model location for studying anthropogenic impacts on biogeochemical cycles in non-pristine mangroves (Vande et al, 2019).

According to the zonation (fringe, basin, impaired area), three mangrove species comprise the vegetation cover in Estero Pargo: at the fringe (approximately 20-30 m wide) along the edge of the tidal channel, the dominant species is the red mangrove (*Rhizophora mangle*), with a few individuals of white mangrove (*Laguncularia racemosa*) also present. Moving to the basin (about 300 m wide), situated behind the fringe, the dominant species shifts to the black mangrove (*Avicennia germinans*). Lastly, there exists a patch (approximately 2.8 ha) of an impaired mangrove area where most of the trees have died, yet a few individuals of *A. germinans* persist (Supplementary figure 1).

The annual average rainfall in the study area is approximately 1,680 mm, with a mean annual temperature of 27 °C (Coronado-Molina et al., 2012). The tidal regime is about 0.33 m (Contreras Ruiz Esparza, Douillet & Zavala-Hidalgo, 2014). There are two periods of minimum (February to August) and maximum (September to January) flooding. The soil bulk density is 0.22±0.008 g cm^-3^ to 0.44± 0.02 g cm^-3^ and organic matter is 8.13 ±1.86% to 12.0 ±3% (Pérez-Ceballos et al., 2022).

### Sampling

To characterize the bacterial community in the sediments of Estero Pargo, we conducted sampling along a conservation gradient, ranging from the most conserved zone, hereafter referred to as non-degraded (ND) or the fringe, followed by the moderately degraded (MD) site or the basin, to the most damaged area, referred to as further degraded (D) or the impaired mangrove. The ND site exhibited no evidence of natural or anthropic factors impacting the forest structure, with most trees alive and the forest covering this zone completely, serving as our reference site. In the MD site, although most trees were alive, signs of degradation were observed as several trees had died. Lastly, the D site experienced further tree mortality, with only a few living trees remaining (Supplementary figure 1).

We established three sampling plots, each measuring 100 m^2^, parallel to the tidal channel within each sampling zone. At the midpoint of each plot, we collected sediment cores with a depth of 50 cm from the soil surface. Subsequently, we subsampled each core at intervals of 0-10 cm, 10-30 cm, and 30-50 cm (hereafter referred to as horizon levels) to obtain samples from the central part of each subsection, corresponding to 5 cm, 20 cm, and 40 cm horizon levels. To address seasonal variations, we collected three replicates from each site during two distinct seasons: the flooding season (January) and the dry season (May) in 2018. All samples were transported to the laboratory and stored at -20°C until processing.

### DNA extraction

After thawing and fully homogenizing the samples, we extracted DNA for molecular analyses using 250 mg of sediment. To ensure a comprehensive representation of the bacterial community at each horizon level, we pooled five replicates of DNA extraction per core section for analysis. We purified DNA from 54 sediment samples according to the manufacturer’s instructions using the E.Z.N.A Soil DNA kit (Omega Biotech, Inc. Georgia, USA).

### Amplification and Sequencing

Genetic libraries were prepared using the primers 341F (CCTACGGGNGGCWGCAG) and 785R (GACTACHVGGGTATCTAATCC) to amplify fragments of the V3-V4 regions of the 16S RNA gene (Ma et al., 2021). DNA samples were prepared for targeted sequencing with the Quick-16S™ NGS Library Prep Kit (Zymo Research, Irvine, CA). The sequencing library was prepared using real-time PCR machines to control cycles and limit PCR chimera formation. All PCR reactions were set to a final volume of 20 uL and treated with the following thermal protocol: 10 min at 95°C for denaturation, then 20 cycles of 95°C for 30 sec, 55°C for 30 sec, and 72°C for 3 min. The final PCR products were cooled at 4°C prior to quantification with qPCR fluorescence readings and pooled together based on equal molarity. The final pooled library was cleaned up with the Select-a-Size DNA Clean & Concentrator™ (Zymo Research, Irvine, CA), then quantified with TapeStation® (Agilent Technologies, Santa Clara, CA) and Qubit® (Thermo Fisher Scientific, Waltham, WA). We used the ZymoBIOMICS® Microbial Community Standard (Zymo Research, Irvine, CA) as a positive control for each DNA extraction. The final library was sequenced on Illumina® MiSeq™ with a v3 reagent kit (600 cycles), and performed with >10% PhiX spike-in.

### Data processing and taxonomy assignment

The quality of the raw reads was analyzed using FastQC (v.0.12.0) (Andrews S, 2010). Subsequently, TrimGalore (v.0.6.10) (Krueger et al., 2023) was used to remove Illumina Universal Adapter (AGATCGGAAGAGC) from the reads. We utilized R (v4.2.2) (R Core Team., 2023) and DADA2 (v.1.16) (Callahan et al., 2016) to filter the reads with the following parameters: the reads were truncated to 290 bases for forward and 200 bases for the reverse. This means that reads longer than these were not allowed. The maximum expected errors (maxEE) which refers to the limit of errors in a sequence at a base level allowed per read, was set at 5 errors per read. Reads with an expected error exceeding this value were excluded. The ambiguous bases (N) were not allowed. For taxonomy assignment, we employed the function assignTaxonomy from the package DADA2 (v.1.16) (Callahan et al., 2016) with the database GreenGenes2 (v.2022.10) (McDonald et al., 2023).

### Phylogenetic reconstruction

We utilized the AlignSeqs function from the DECIPHER package (Wright, 2017) to align the 16S RNA sequences. Subsequently, we employed the dist.ml function from Phangorn (Schliep, 2011) to generate a distance matrix using the F81 substitution model. We then employed the nj function to construct an unrooted tree using Neighbor Joining.

### Community analysis

We explored the microbial community accounting for horizon depth, zone and season variables. Visualization of each parameter option was achieved through non-metric multidimensional scaling (NMDS) analysis, employing Bray-Curtis distance. Based on the exploratory analysis, the sample zr2502_16_R1 which could potentially have skewed the analysis, was excluded (Supplementary Figure 2). Subsequently, our dataset was filtered to encompass only those amplicon sequence variants (ASVs) present in at least 10% of the samples.

We employed the plot_richness function from phyloseq (McMurdie & Holmes, 2015) to investigate alpha diversity patterns using the Chao1, Simpson, and Shannon indices. Significance testing was conducted through a Wilcoxon test, with adjustments for multiple comparisons using the Bonferroni method. Additionally, we employed ANOVA followed by Tukey’s test for further analysis. We employed microViz (v.0.12.1) (Barnett, Arts & Penders, 2021), a comprehensive package to conduct both Principal Component Analysis (PCA) and Principal Coordinate Analysis (PCoA) to elucidate the underlying patterns of variation in our dataset. Prior to analysis, the data underwent a centered log-ratio (clr) transformation for PCA, enabling effective exploration of compositional differences while mitigating the impact of spurious correlations. For PCoA, we did not apply any transformation, allowing for a direct examination of dissimilarity among samples using Bray Curtis distance and UniFrac distance. We corroborate the observed patterns performing a PERMANOVA test. These analyses collectively provided insight into the structure and distribution of the data, facilitating the interpretation of ecological dynamics.

Differentially abundant bacteria were detected using the microbiomeMarker package (v.1.8) (Cao, 2021), specifically by employing the run_aldex function with the glm_anova method, assessing all taxonomic ranks. This analysis was conducted across all data pairs, resulting in a list of differentially abundant taxa, which is available in Supplementary Table 1.

### Metabolic inference

PICRUSt2 was used for the prediction of the functional abundances (Douglas et al., 2020), where ASVs are placed into a reference tree with reference genomes; hidden-state prediction approaches are used to infer the genomic content of sampled sequences using several databases such as enzymes (ECs), KEGG orthologues (KOs), cluster of orthologous genes (COGs) and metabolic pathways (MetaCyc). To obtain and visualize differentially abundant data (KOs), *ggpicrust2* package (v.1.7.3) in R was used (Yang et al., 2023). The method applied to asses differentially abundant data was ALDEx2 (Fernandes et al., 2013)

## Results

### Microbial diversity dynamics across mangrove ecosystem degradation levels

Our study aimed to elucidate how microbial communities respond to various levels of mangrove ecosystem degradation, classified as non-degraded (ND), moderate degradation (MD), and degraded (D). Additionally, we accounted for seasonal variations and sampled three different depths (5, 20, and 40 cm) as part of our experimental design (see Figure 1A). We conducted a comprehensive analysis of microbial communities within the mangrove ecosystem, utilizing 619,326 raw sequence reads, which led to the identification of 11,469 ASVs. Subsequently, we filtered out low-abundance taxa (present in less than 10% of the samples), retaining 648 ASVs for further analysis. We hypothesized that increasing levels of mangrove ecosystem degradation would lead to a decline in microbial diversity, accompanied by shifts in microbial community structure influenced by both season and depth (Figure 1B). To assess this hypothesis, we analyzed alpha diversity patterns. The Chao1 index, revealed significant differences between MD and D sites, where the MD site is more diverse. We conducted pairwise comparisons using the Wilcoxon rank sum test with continuity correction to assess differences in Chao1 richness index among sites. The analysis showed a statistically significant difference between the MD and D sites (Wilcoxon rank test, p = 0.017), while no significant difference was observed between the ND and D sites. P-values were adjusted for multiple comparisons using the Bonferroni method. Consistent with the Chao1 index findings, the Shannon diversity index showed a noticeable decrease in diversity as degradation levels increased, which was significant among MD and D. However, the Simpson diversity index did not reveal significant differences between degradation sites; however, it indicated elevated values, suggesting the dominance of certain microbial groups (Figure 1C).

**Figure 1.**
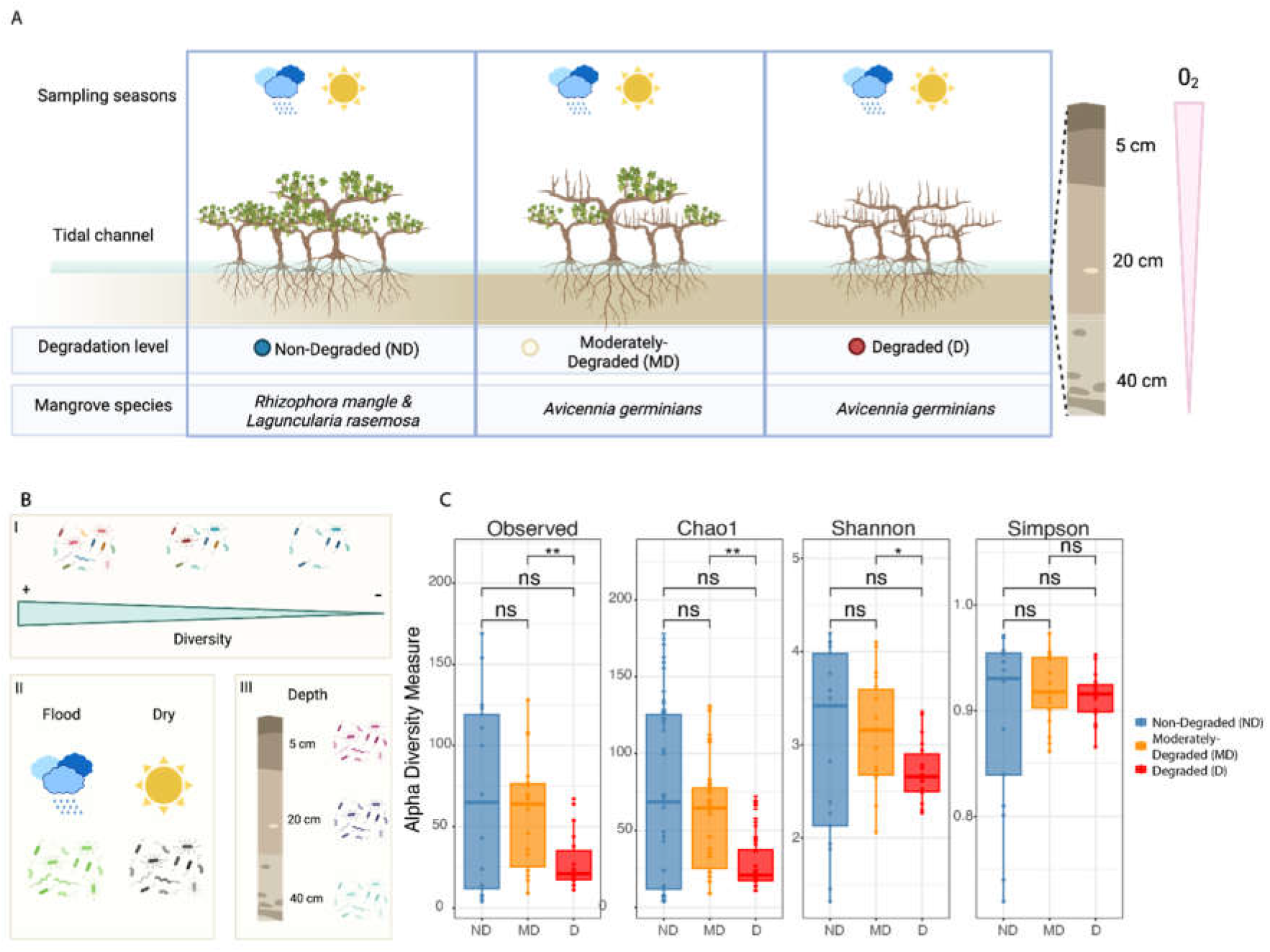
Overview of Experimental Design and Alpha Diversity. A) Schematic representation of the experimental design depicting three distinct mangrove sites, each characterized by specific mangrove species. B) Hypothesized alterations in microbial diversity: anticipation of a reduction in microbial diversity (I), fluctuations based on seasonal variations (II), and shifts relative to depth (III). C) Comparative analysis of alpha diversity, employing three distinct indices, across degradation sites, significance of the comparison * = 0.05, ** = 0.01 and ns = not significant.

To comprehend alpha diversity patterns, we divided the dataset by depth and assessed diversity across seasons. Employing the Simpson index to quantify species dominance and the Shannon index to consider species richness and evenness, we obtained a comprehensive view of community diversity. Notably, the dry season consistently displayed higher diversity than the flood season, supported by ANOVA and Tukey tests revealing significant seasonal differences (p-adjust = 0.007). Diversity at the 5 cm depth was linked to water loss, with diversity declining during the dry season according to degradation levels, contrasting with the flood season. Statistical analysis confirmed significant mean differences between non-degraded and degraded sites for the Shannon diversity index (p-adjust = 0.011) and Simpson index (p-adjust = 0.023; Figure 2). At the 20 cm depth, diversity patterns were similar across seasons, with degradation leading to decreased diversity, though not statistically significant. Nonetheless, the Simpson index highlighted higher dominance during the dry season, indicating prevalent groups during this period. At the deepest horizon of 40 cm, the dry season appeared substantially more diverse than the flood season, possibly due to water presence altering oxygen gradients, leading to collapse of anoxic microorganisms during normal conditions. Statistical tests revealed significant diversity differences between degraded and non-degraded sites (p-adjust = 0.060) using the Shannon index. However, in the flood season, no significant differences were found using the Shannon index. Nevertheless, significant differences were noted between non-degraded and moderately degraded sites (p-adjust = 0.01), as well as between non-degraded and degraded sites (p-adjust = 0.009) using the Simpson index.

**Figure 2.**
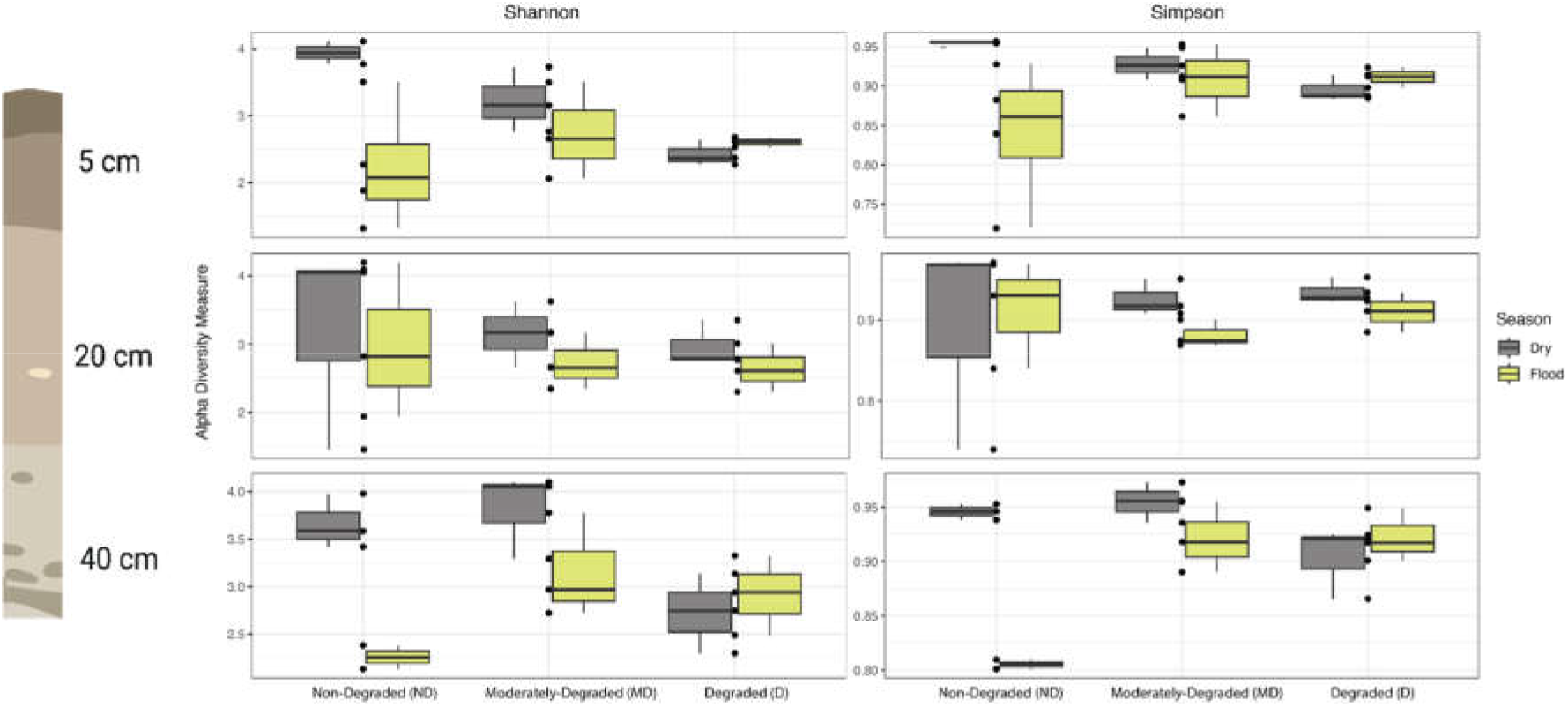
Alpha diversity across degradation sites considering season and depth. The Shannon and Simpson indices depict diversity patterns, highlighting variations across different degradation sites.

### Depth and zone drive changes in microbial community structure

To explore the microbial community dynamics, we used the clr-transformation and Principal Component Analysis (PCA), we delineated distinct microbial communities between ND and MD sites, particularly noticeable during the dry season in the MD site (Figure 3A). Principal Coordinate Analysis (PCoA) on the untransformed data further revealed a distinct clustering associated with the ND site (Figure 3B), suggesting that there are characteristic microbial groups in this condition. Moreover, depth analysis via PCoA’s third dimension demonstrated a consistent microbial community at the 40 cm horizon across degradation levels and seasons (Figure 3C), contrasting with upper horizons where degradation level (Figure 3D), especially in the dry season (Rectangles in Figure 3D), significantly influences microbial composition. PERMANOVA analysis conducted with the adonis method reveal significant effects of the three environmental variables zone, season and depth (p = 0.0001), however based on the R-squared value we observed that depth and zone explain large proportion of variation (0.07199 and 0.07264, respectively), while season only explained the 0.03531 (Supplementary Table 2).

**Figure 3.**
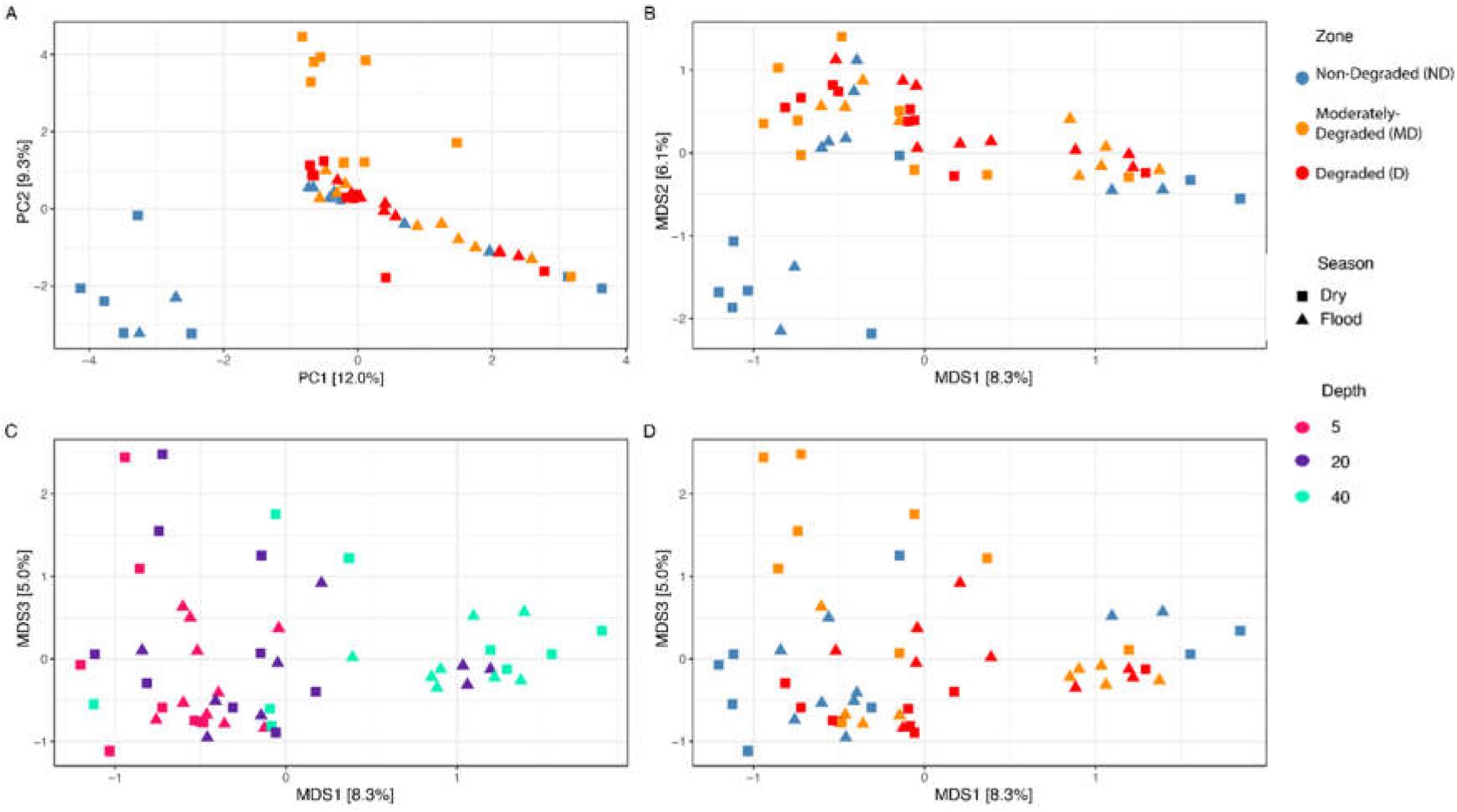
Sample Distribution Based on Environmental Variables. A) Principal Component Analysis illustrating sample distribution by zone and season, differentiated by color and shape, respectively. B) Principal Coordinate Analysis showcasing the first and second dimensions. C and D) First and third dimensions of Principal Coordinate Analysis. Panel C emphasizes depth through color, while D highlights the zone.

We examined the microbial community composition using UniFrac distance, which measures genetic distance and highlights differences in microbial communities based on lineage composition. Our analysis, employing a principal coordinate analysis (PCoA) and PERMANOVA, demonstrated that our three variables play significant roles in shaping the community structure (P < 0.001; see Supplementary Table 3). Regarding depth, microbial lineages from the 40 cm horizon stand out from the rest of the community. Additionally, they appear to be closely associated with the flood season (Figure 4A,C). Meanwhile, our findings suggest that the degraded zone harbors a distinct composition of lineages compared to other degradation stages (Figure 4B).

**Figure 4.**
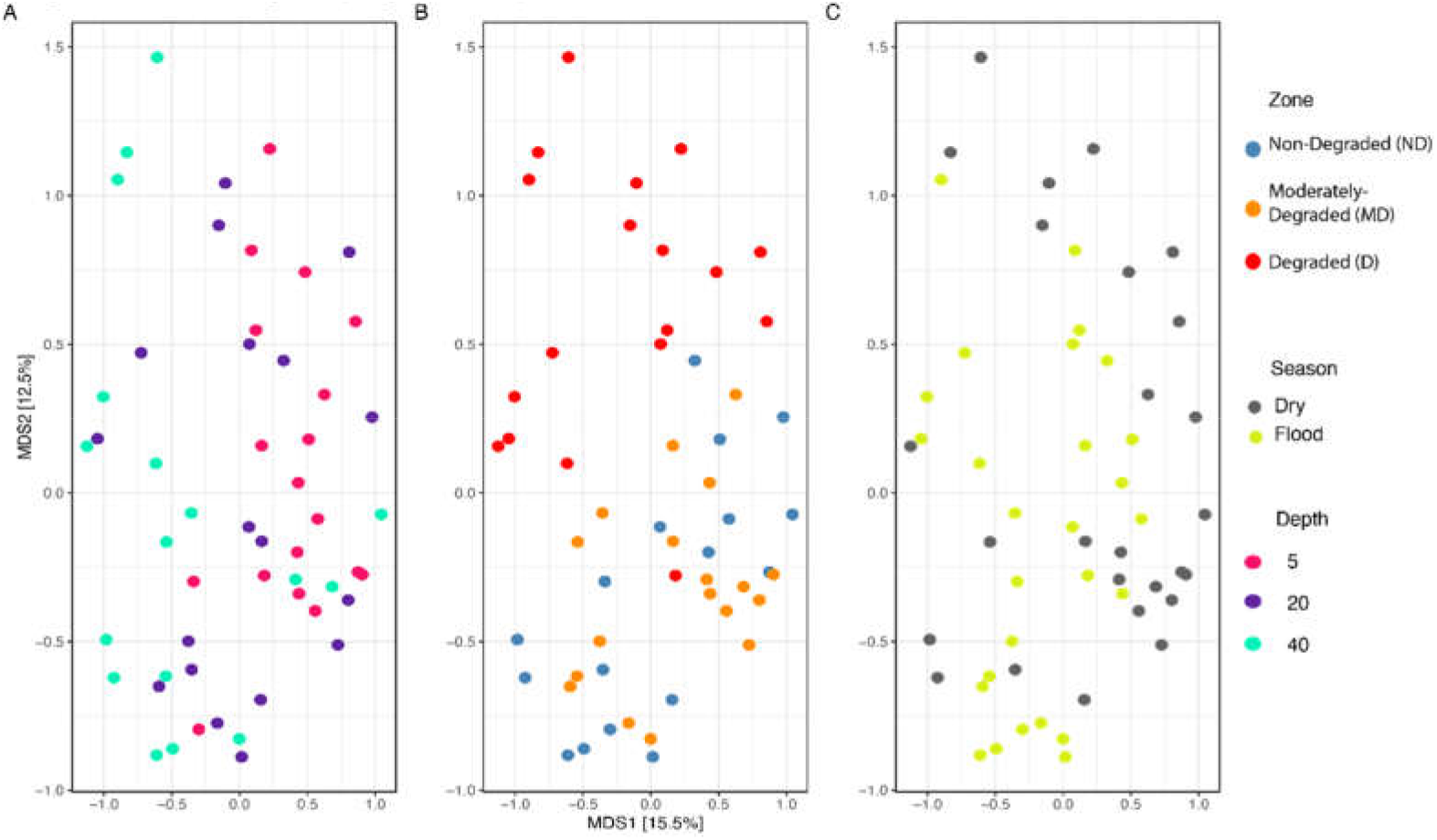
Microbial community structure depicted using UniFrac distances. Each panel represents a different variable: A) Depth, B) Zone, and C) Season, color-coded accordingly.

### Shifts in Gammaproteobacteria and Firmicutes

To elucidate the microbial diversity contributing to differences in community structure, we reconstructed a phylogenetic tree. This analysis unveiled distinct patterns among Gammaproteobacteria and Firmicutes across degradation sites, with Gammaproteobacteria prevalent in both ND and MD sites, while Firmicutes exhibited greater diversity and spread across all degradation stages (Figure 5). Notably, within Proteobacteria, particularly the Vibrionaceae family, we observed heightened prevalence in both ND and MD sites.

**Figure 5.**
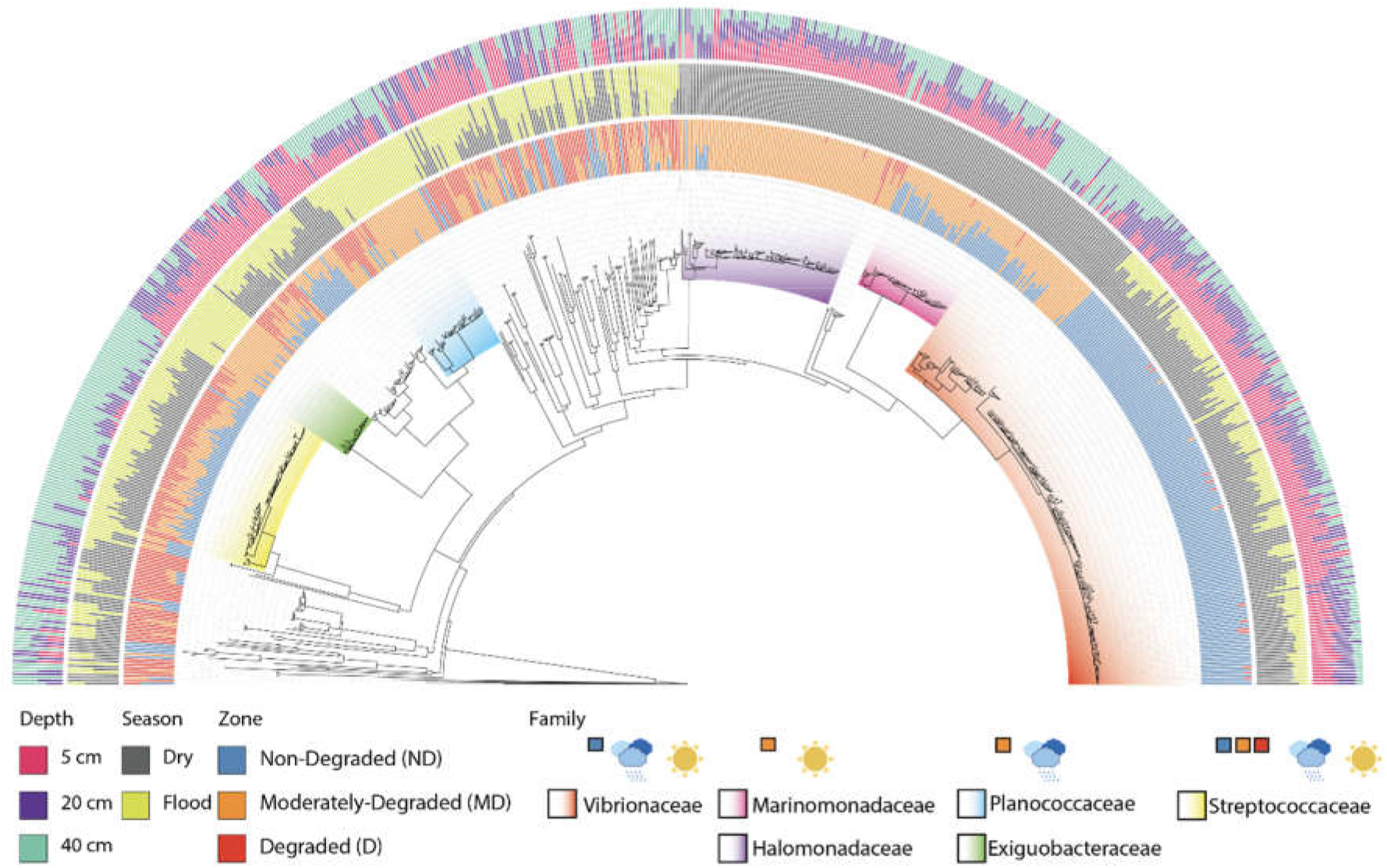
The phylogenetic reconstruction of 16S rRNA Amplicon Single Variant (ASVs) present in Estero Pargo. The inner circle delineates the sampling zone, season, and depth. Each bar indicates the abundance of the corresponding ASV. Clade color indicates taxonomy at the family level. Above the names we highlighted the zone and season from which those families were found.

However, a subset of Vibrionaceae was exclusively confined to the ND site, contrasting sharply with the dominance of Marinomonadaceae and Halomonadaceae in the MD. Additionally, specific lineages of Nitrospirota were solely identified in the D site, suggesting a potential association with our observed alpha diversity patterns that show high dominance. Conversely, within the Firmicutes phylum, clusters associated with *Lactococcus* from the family Streptococcaceae were uniformly distributed across all sites, indicating a significant role in the ecosystem. Regarding seasonal dynamics, Exiguobacteraceae and Planococcaceae were predominantly concentrated in MD sites. Interestingly, the prevalence of Marinomonadaceae and Halomonadaceae in MD sites correlated with the dry season, while Exiguobacteraceae and Planococcaceae thrived during the flood season. Furthermore, our spatial analysis revealed depth-related distribution patterns, particularly pronounced in ND and MD sites. Gammaproteobacteria, linked with Marinomonadaceae, Halomonadaceae, and Vibrionaceae, were predominantly found at the surface (5 cm depth), whereas Firmicutes associated with Lactococcus were notably prevalent at 40 cm depth, spanning all zones and seasons. This observation suggests a central role for Lactococcus-related Firmicutes in ecosystem dynamics.

We utilized the analysis of differential abundance considering sample and scale variations (ALDEx2; see Supplementary Table 1) to examine disparities in abundance across diverse environmental parameters. Among sites categorized as ND and MD, certain taxonomic groups emerged as distinctive to the ND site such as the phyla Desulfobacterota and Chloroflexota, along with the ASVs classified under the species *Vibrio rumoiensis*, as previously observed in the phylogenetic tree (Figure 4). When contrasting MD with D site, lineages from Firmicutes such as Rossellomorea and Lactococcus predominated, alongside Halomonadaceae, indicative of the MD site. Additionally, lineages from the phylum Nitrospirota, specifically from the class Thermodesulfovibrionia, distinguished the D site. Seasonal dynamics revealed a pronounced pattern of higher abundance of lineages from the families Marinomonadaceae and Halomonadaceae during the dry season (Figure 5), correlating with the MD site. On the contrary, during the flood season, Firmicutes and Desulfobacteriota were predominant. Depth-related variations were notable, particularly between the surface (5 cm) and deeper horizons (20 cm and 40 cm). Firmicutes, Actinobacteriota, and Desulfobacteriota were predominant at the surface, while Nitrospirota, especially Thermodesulfovibriota, and Pseudomonadaceae were prevalent at 40 cm.

### Season and zone influence on metabolism patterns

Phylogenetic placement of ASVs in a genome tree revealed that the dry season exhibited higher diverse presence of enzymes (ECs), metabolic pathways, KEGG orthologues (KOs) and COGs compared to the flood season (Figure 6) which is in agreement with what we observed regarding taxonomic diversity (Figure 2). The differential analysis of KO’s abundance indicated a significantly greater statistical difference in several types of metabolism such as amino acid metabolism and the degradation of xenobiotic compounds (Figure 6B and C) in the non-degraded sites compared to the degraded and moderately degraded ones.

**Figure 6.**
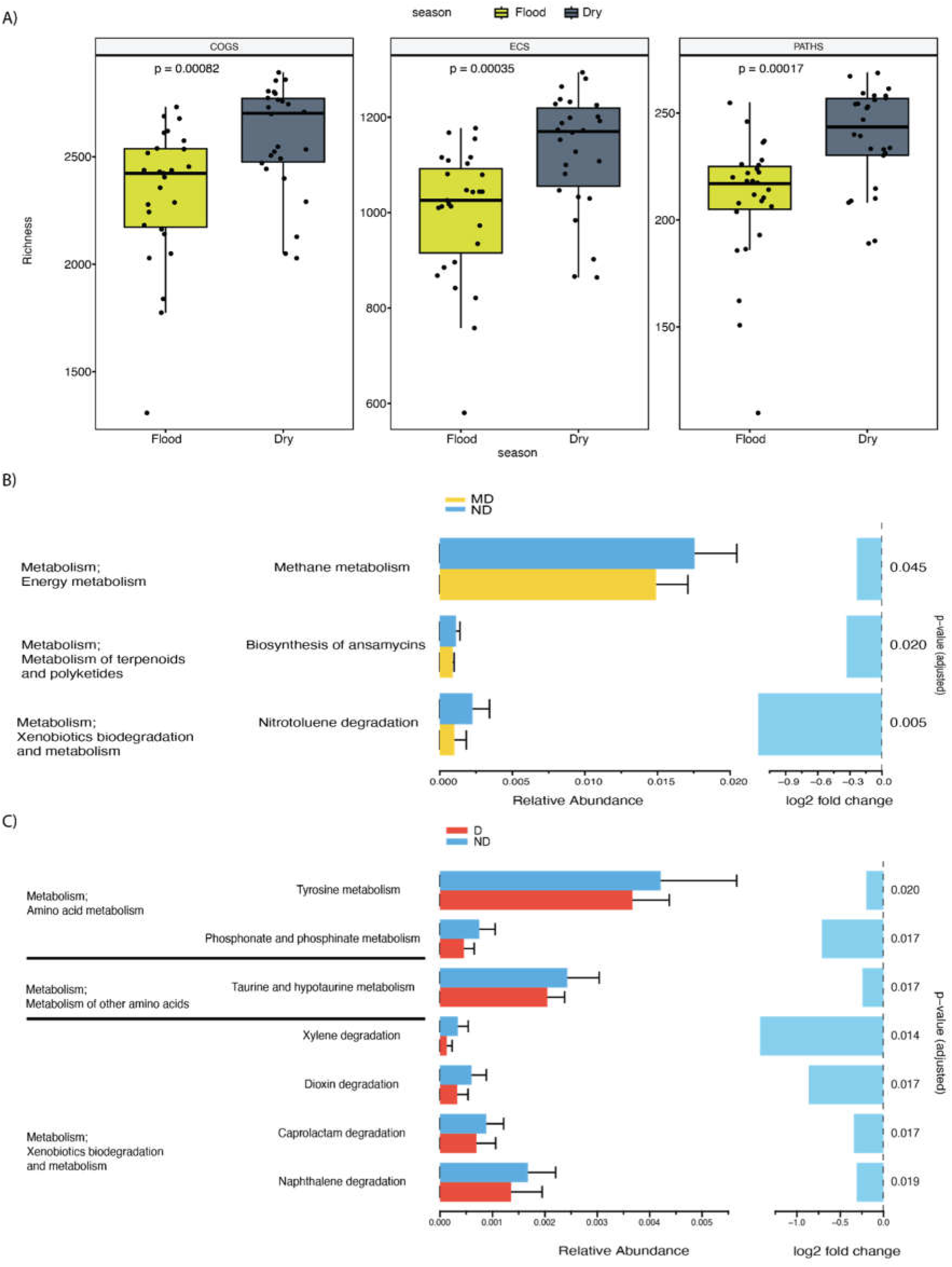
Metabolism overview. A) Abundance of predicted enzymes categorized as COG, EC, KO and Pathways. Differential abundance analysis (ALDEx2) in KO’s accounting for the different zones: B) MD vs ND and C) D vs ND.

## Discussion

Our study aimed to explore how microbial communities respond to varying degrees of degradation within mangrove ecosystems, characterized by altered hydrological connectivity and increasing urbanization. Our findings show the dynamic nature of surface sediment layers, which exhibit heightened susceptibility to change. This was particularly evident at the non-degraded and moderately degraded sites. At these sites, distinct microbial compositions emerged, with one site dominated by Vibrionaceae and the other by bacteria like Marinomonadaceae and Halomonadaceae, displaying clear seasonal patterns. Despite this, certain bacteria persist in deeper sediment layers, potentially playing key roles as early colonizers in ecosystem restoration efforts. Furthermore, our observations highlight the impact of water flow on microbial diversity, being the dry season more diverse than the flood.

As anticipated, we noted a decrease in microbial diversity correlated with the degradation level. However, we found no reduction in functional diversity according to the degradation level. This led us to propose that the key molecular functions may be conserved through the loss of hydrological connectivity. This could be understood by the term of functional redundancy, which refers to the coexistence of multiple distinct taxa or genomes capable of performing the same focal biochemical function (Louca et al., 2018), facilitating the maintenance of ecosystem functions despite a loss in taxonomic diversity (Li et al. 2021; Haroon et al. 2013; Waite et al. 2020; Wang et al. 2023). However, regarding seasonality, the dry season consistently exhibited higher diversity compared to the flooding season, with a parallel pattern observed in metabolic diversity; this pattern may be related to the dominance of microbial groups specifically related to Proteobacteria. For example, bacteria residing in mangrove sediments play a pivotal role in denitrification, the process through which they convert nitrate (NO_3_^-^) and nitrite (NO_2_^-^) into nitrogen gas (N_2_). Various bacterial taxa, such as Pseudomonas, Paracoccus, and Bacillus, contribute to this essential process at different sediment depths within mangrove ecosystems (Haroon et al., 2013). Similarly, sulfate-reducing bacteria (SRB) like Desulfobacter, Desulfovibrio, and Desulfuromonas are crucial in mangrove sediments, where they convert sulfate (SO_4_^2-^) into sulfide (S^2-^). These SRBs can be found at various sediment depths, actively participating in sulfate reduction processes (Waite et al., 2020). Moreover, bacteria involved in carbon metabolism, such as Clostridium, Cellulomonas, and Acetobacterium, contribute to cellulose degradation and fermentation processes (Wang et al., 2023) within mangrove sediments. These bacteria also exhibit functional redundancy across sediment depths, indicating their importance in maintaining ecosystem function. Furthermore, their distribution is influenced by spatial contiguity and reciprocating tidal flows (Li et al., 2021). We are aware of the bias that metabolic prediction through 16S RNA comes with, however, we believe it gives us some general patterns regarding what we can expect and help us to raise new hypotheses regarding the metabolic potential in the community structure.

In mangrove ecosystems, the interplay of water dynamics plays a crucial role in shaping microbial communities, with significant implications for their diversity and composition (Luis et al., 2019). During the flooding season, water flow fluctuations influence the transition between aerobic and anaerobic conditions, thereby impacting microbial diversity. Observations at different sediment horizons revealed interesting patterns: at 5 cm depth, diversity tends to increase with degradation levels, possibly due to input from external water sources like the ocean or lagoon. A possible source of diversity variation is the tidal importation of organic matter. An increase in bacterial diversity has been observed in the top layers of mangrove sediments due to the input of organic matter that tidal regimes allow, creating microhabitats that increase the diversity of heterotrophic bacteria (Zhu et al. 2018). Similarly, at 40 cm, diversity follows a similar trend, yet notably lower diversity at the ND site suggests limitations imposed by anaerobic conditions induced by water saturation, which likely restrict the presence of aerobic-obligate bacteria. Hence, variation in hydrology emerges as a key driver shaping microbial communities in these ecosystems. In the dry season, when water flow ceases, bacterial diversity undergoes significant transformations depending on the duration of isolation from external water sources. Prolonged isolation leads to pronounced shifts in microbial diversity. However, when floods reintroduce water flow, microbial diversity decreases as new species from external sources dominate the environment. Similar to our findings, other studies demonstrated that in freshwater systems, the bacterial communities respond to dry and wet seasons by changing their functional and taxonomic diversity, showing an increased alpha diversity during the dry season (Ren et al., 2019). Such results can be associated with higher nutrient concentrations, lower water dilution capacity, and reduced degradation of organic matter (Ren et al. 2019). The availability and susceptibility to change of dissolved organic matter are critical factors influencing bacterial community composition, particularly in upper soil horizons. Understanding these dynamics provides insights into the intricate relationships between water dynamics and microbial communities in mangrove ecosystems.

Due to the intermittent flooding caused by tides in mangroves, the availability of nutrients and the concentration of oxygen and salinity, among other factors, vary significantly (Basak et al., 2016). It is known that the availability of nutrients determines the structure of the microbial communities. For example, S concentration and anoxic conditions can increase the abundance of Deltaproteobacteria and Epsilonproteobacteria as organisms representative of these phyla reduce or oxidize S compounds (Lin et al., 2019). It is known that the percentage of organic matter, pH, and seasonal water saturation influence the composition and structure of the microbial community and the abundance of specific functional groups (Ansola, Arroyo & Sáenz de Miera, 2014; Arroyo, Sáenz de Miera & Ansola, 2015; Ding et al., 2015).

Therefore, seasonal variation in hydrological connectivity, water flow, or stagnation, mainly at site D, influenced the structure and composition of the bacterial community, where we observed an increase in the abundance of Nitrospira, a bacteria that participates in the nitrogen cycle, which was abundant on this site, suggesting that an increase of ammonium exist in this environment (Daims & Wagner, 2018; Meng et al., 2022).

In non-degraded mangrove ecosystems, a higher abundance of KO’s associated with amino acid metabolism reflects optimal environmental conditions. Amino acids serve pivotal roles as precursors for essential biochemical processes including protein synthesis, nucleotide formation, chlorophyll production, and hormone regulation. The significant difference in the relative abundance of these KO’s in the ND compared to MD and D zones, indicates efficient nitrogen utilization for growth and development. Therefore, the greater presence of amino acid metabolism-related KO’s in non-degraded mangroves signifies a balanced and healthy ecosystem state supporting vigorous plant metabolism and ecological stability. (Shah et al., 2021; Ningsih et al., 2020).

On the other hand, our results were similar to those reported for riverine flood plains, where the abundance of anaerobic bacteria increased when the system remained flooded for a more extended period (Argiroff et al., 2017). We observed higher diversity and dominance of aerobic bacteria such as Firmicutes and Proteobacteria in the dry season, which might be signals of a loss in hydrological connectivity and water flow.

Tidal influence may shape bacterial diversity by introducing taxa from external habitats (Zhu et al. 2018). As the ND site was consistently influenced by tidal variation in Estero Pargo, it showed distinctive groups of Vibrionaceae. *Vibrio* species are natural inhabitants of aquatic environments, including sediments, estuaries, and marine coastal waters. They can live in both saline waters with 30-35 ppt and low saline environments (Wong & Griffin, 2018). As well, *Vibrio* species have been found as part of the microbiota of *R. mangle* (Gomes et al., 2014) which may explain its presence in the ND site, which is dominated by this family. Vibrio species isolated from salt marshes have shown the capacity of fixing nitrogen, which suggests that the presence of *Vibrio* in the ND site could be also playing this role (Criminger et al., 2007). And other *Vibrio* directly isolated from *R. mangle* have been shown to be able to inhibit bacteria and fungal phytopathogens (Rameshkumar & Nair, 2009). Recent increases in *Vibrio* abundance have been related to human activities that can negatively affect aquatic or marine habitats (Narayanan et al., 2020). For example, aquaculture is thought to be an activity that can increase the abundance of *Vibrio* species because organisms such as rotifers and Artemia, used as live feed, can be hosts of several *Vibrio* species (Sanches-Fernandes et al., 2022). In addition, high densities of cultured fish can, in turn, increase the abundance of *Vibrio* species that can be released to natural environments when ponds are cleaned and water is discharged to water courses (Sampaio et al., 2022). In our study area, the growing fish farming activities could contribute to the recorded *Vibrio* abundance in Estero Pargo.

At the MD site, we observed a notable shift in microbial diversity. During the dry season, Halomonas and Marinomonas were notably abundant, whereas during the flood season, Exigoubacterium emerged as dominant. Although all three groups of bacteria are recognized as halophilic, changes in salt concentration may favor some lineages over others (Edbeib, Wahab & Huyop, 2016; Chen et al., 2017). Mangrove sediments show oxygen gradients; typically, the upper layers have high oxygen concentrations, while the lower layers have less oxygen and high rates of other elements (sulfur, nitrogen, and methane), suggesting multiple vertical niches. Therefore, the diversity changes at the MD site could reflect these multiple niches. Initially, we speculated that the MD site might resemble a pristine mangrove environment due to the high presence of Vibrionaceae, as in the ND site. However, given the tight correlation between degradation levels and decreased water flow, it is plausible that the MD site reflects how arid environmental conditions act as a selection factor for halotolerant lineages.

*Halomonas* has been found in other mangrove ecosystems as part of the core microbiome, it has been suggested that the production of ectoine an osmolyte molecule might help the host to cope with osmotic stress. Therefore, considering the holobiont hypothesis, this bacteria could be interacting with other organisms within the ecosystem, probably the mangrove tree to provide this type of protection (Wainwright et al., 2023). On the other hand, Marinomonas has been shown to participate in the conversion of dimethylsulfoniopropionate (DMSP) to dimethyl sulfide (DMS) which has been shown in corals that they help them in the sulfur cycle, therefore we believe that could be also happening here (Wainwright et al., 2023).

Our study revealed enrichment of halophilic bacteria in certain conditions, this suggests the potential use of some halophilic bacteria to protect the plants from saline stress as it has been shown before (Bharti et al., 2015).

The MD site dominated by *Avicieana germinias* showed a different microbial composition compared to what has been described previously, Desulfatiglans has been described as an abundant lineage associated to this type of tree, however in our study we did not find this to be any abundant, similar thing happened with Ignavibacteriales, Phaeodactylibactes and SAR324, which were associates to *A. germinis* (Gómez-Acata et al., 2023). This could be related to 1) we sampled a human-impacted ecosystem. Therefore, that could change the microbial structure compared to other studies; 2) differences in substrate composition, which in previous studies is karstic; and 3) our low sequence coverage, which may mislead our conclusions. However, this also shows the uniqueness of microbial communities associated with specific trees.

Despite the higher variability in species diversity observed in the upper sediment horizons, we observed that the diversity at 40 cm was preserved in all the degradation levels. This result suggests that environmental conditions stay stable at greater depths for longer, allowing the bacterial community to remain unchanged over more extended periods. We believe this community could be used in restoration efforts as a seed community to enhance the process. Interestingly, Lactococcus was observed as a dominant lineage at 40 cm, this lineage is usually constrained to habitats like food related, however some strains have been isolated from mangrove ecosystems and their production of bacteriocin which are molecules secreted to prevent the growth of other bacteria, has been used in other plant systems to prevent microbial growth (Hwanhlem et al., 2013; Kleerebezem et al., 2020).

Further research is needed to fully understand the metabolic potential underlying our hypothesis. We are eager to explore a metagenomic approach to testing our theories, delving deeper into the microbial communities involved. Additionally, we aim to conduct culturing experiments focusing on specific bacterial strains we believe could play a crucial role in ecological restoration efforts. This multidimensional strategy will not only enrich our understanding of the ecosystem dynamics but also pave the way for practical applications in environmental conservation.

## Conclusion

Here we investigated the impact of lost of hydrological connectivity in the microbial communities of a mangrove ecosystem situated at Ciudad del Carmen, Campeche in the Gulf of Mexico, we found that this lost of connectivity at first in the moderate degraded site is accompanied by the selection of key microbes that can cope with dry and highly halophilic conditions such as Halomonas, Marinomonas and Exiguobacterium, however, there were patterns associated with seasons. The upper levels of the sediments are highly influenced by human activities and shape microbial communities, however deeper horizons, especially during the dry season are highly diverse and can be preserved longer. We believe this community could be used in restoration efforts as a seed community to enhance the process of a new mangrove forest restoration.

## Author Contributions

M.V.R.L, R.P.C, A.Z.J, and J.C.C.D conceived the study and analysis. R.P.C, A.Z.J, A.P.D, O.C.H, and J.C.C.D conducted field work and DNA extraction. M.V.R.L and J.C.C.D wrote the first draft of the manuscript. M.V.R.L and L.D.P.G performed bioinformatic analysis. S.E.H.P performed the metabolic prediction analysis. M.V.R.L prepared figures and tables. R.P.C, A.Z.J, and J.C.C.D got funding. All authors contributed and approved the final version of the manuscript.

## DNA Deposition

Amplicon sequences are publicly available through the MG-RAST project entitled Estero Pargo under the link: https://www.mg-rast.org/linkin.cgi?project=mgp104729.

## Acknowledgements

We thank Josefina Santos Ramírez, Tomás Zaldívar Jiménez, Ricardo Ortegón Herrera, Mario Alejandro Gómez Ponce, Hernán Álvarez Guillén, Andrés Reda Deara their assistance with logistics and field data collection; added a Citlalli Garrido Abreu for their lab analysis.

