## Supplementary figures and images for "Flooding patterns shape microbial community in mangrove sediments"

### Supplementary figure 1

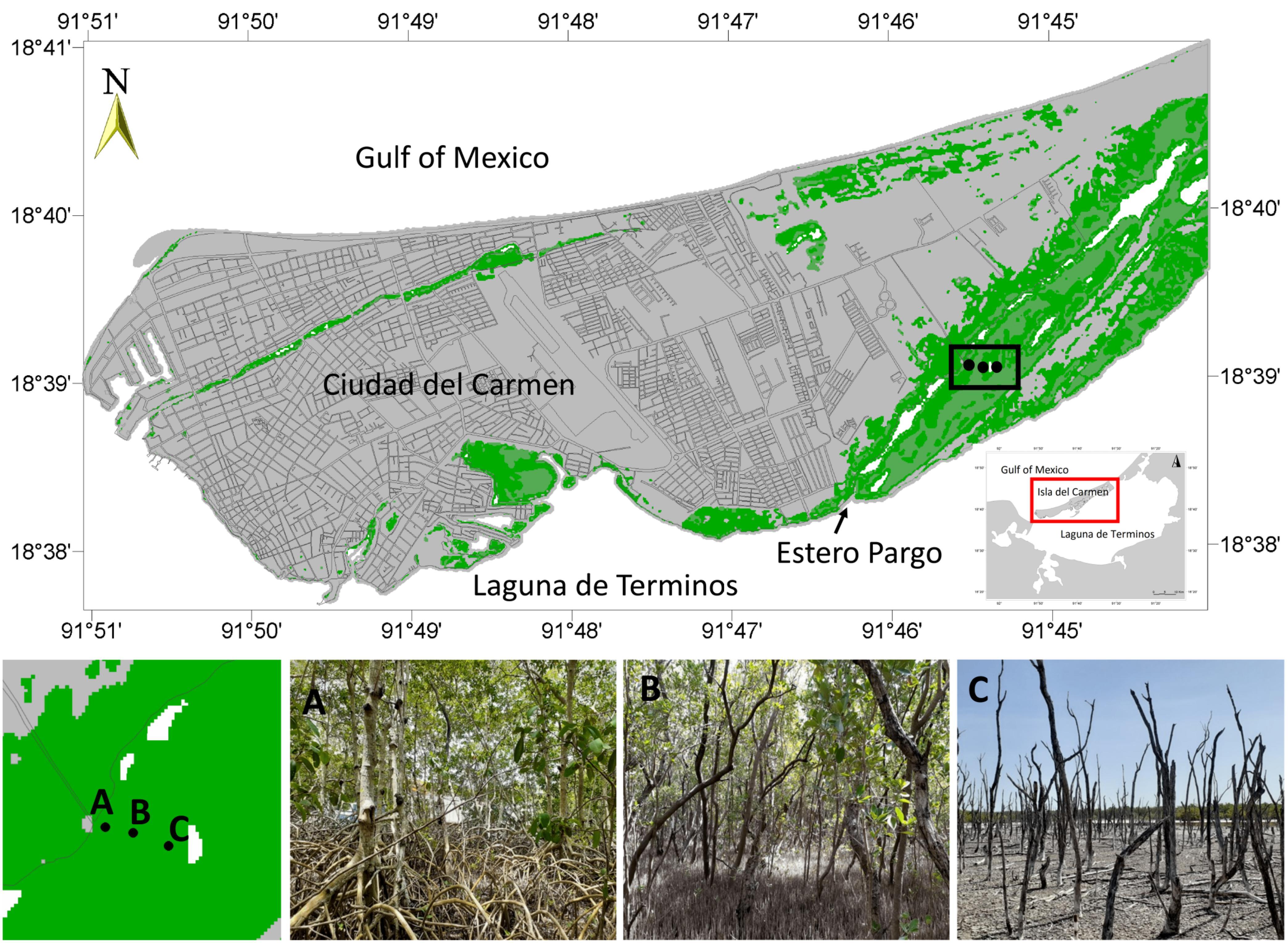

### Supplementary Figure 2

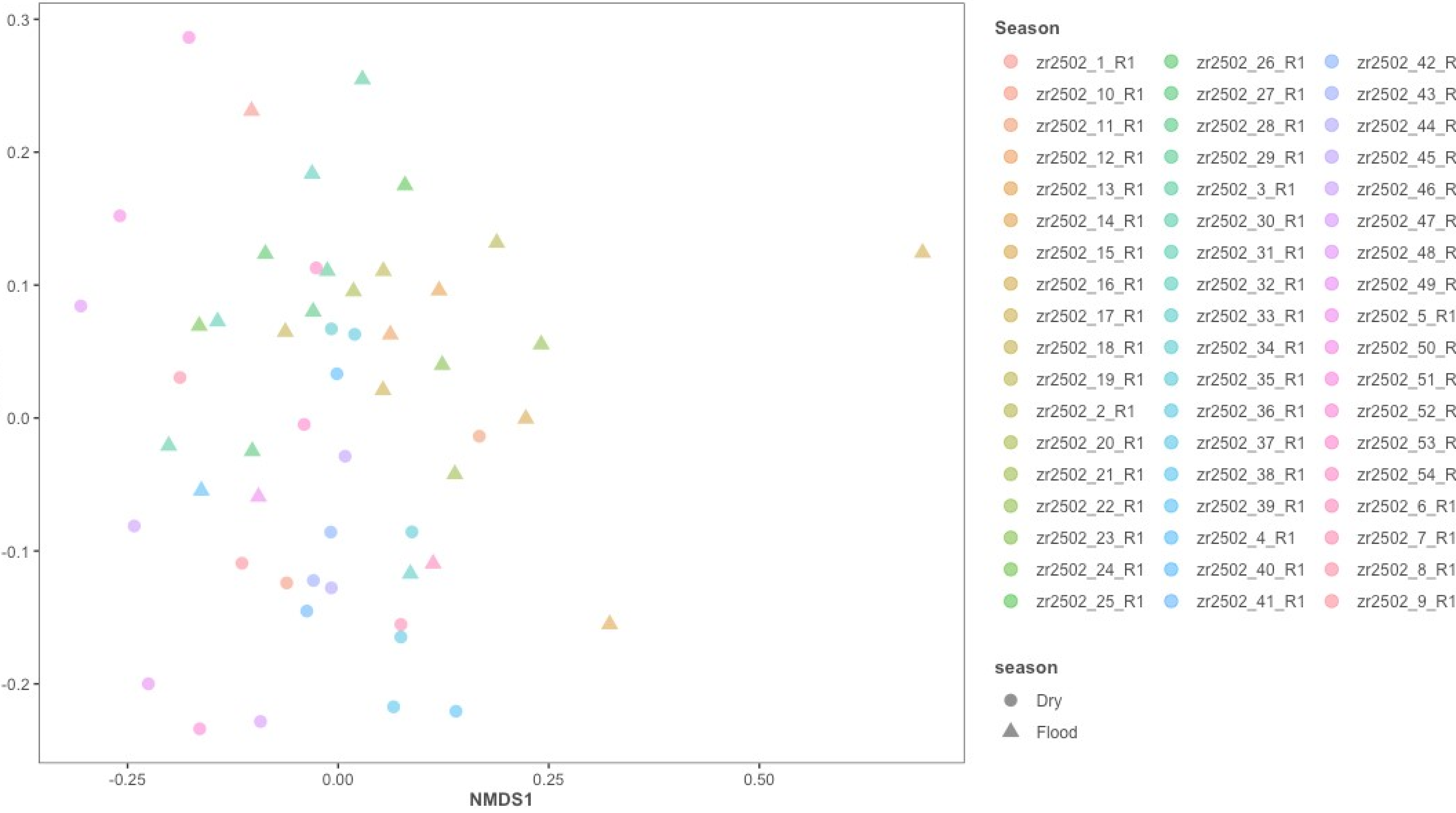
