## Supplementary Table 1 for "Flooding patterns shape microbial community in mangrove sediments"

**Supplementary Table 1. Differential abundance analysis**

| feature | enrich_group | ef_F_statistic | pvalue | padj | comparacion |
|---|---|---|---|---|---|
| d__Bacteria\|p__Desulfobacterota_I\|c__Desulfobacteria\|o__Desulfobacterales | 5 | 11.084649 | 0.0075291 | 0.0075291 | 5-20 |
| d__Bacteria\|p__Desulfobacterota_I\|c__Desulfobacteria | 5 | 8.1564819 | 0.01840262 | 0.01840262 | 5-20 |
| d__Bacteria\|p__Desulfobacterota_I\|c__Desulfobacteria\|o__Desulfobacterales\|f__SG8-13 | 5 | 9.07159493 | 0.01958996 | 0.01958996 | 5-20 |
| d__Bacteria\|p__Desulfobacterota_I\|c__Desulfobacteria\|o__Desulfobacterales\|f__SG8-13\|g__SG8-13 | 5 | 8.88552424 | 0.02216601 | 0.02216601 | 5-20 |
| d__Bacteria\|p__Desulfobacterota_I\|c__Desulfobacteria\|o__Desulfobacterales\|f__SG8-13\|g__SG8-13\|s__sp001303025 | 5 | 8.05118388 | 0.02484219 | 0.02484219 | 5-20 |
| d__Bacteria\|p__Actinobacteriota | 5 | 6.08186827 | 0.02986505 | 0.02986505 | 5-20 |
| d__Bacteria\|p__Actinobacteriota | 5 | 20.722733 | 0.0004501 | 0.0004501 | 5-20 |
| d__Bacteria\|p__Nitrospirota_A_437815 | 40 | 13.8634328 | 0.00277244 | 0.00277244 | 5-40 |
| d__Bacteria\|p__Nitrospirota_A_437815\|c__Thermodesulfovibrionia | 40 | 12.9732788 | 0.0045306 | 0.0045306 | 5-40 |
| d__Bacteria\|p__Firmicutes_D\|c__Bacilli\|o__Bacillales_G_286742\|o__Bacillales_G_286742_f__\|o__Bacillales_G_286742_f___g__\|o__Bacillales_G_ | 5 | 11.6125075 | 0.00486331 | 0.00486331 | 5-40 |
| d__Bacteria\|p__Firmicutes_D\|c__Bacilli\|o__Bacillales_G_286742\|o__Bacillales_G_286742_f__\|o__Bacillales_G_286742_f___g__ | 5 | 12.0559676 | 0.00608699 | 0.00608699 | 5-40 |
| d__Bacteria\|p__Firmicutes_D\|c__Bacilli\|o__Bacillales_G_286742 | 5 | 11.7612634 | 0.00635397 | 0.00635397 | 5-40 |
| d__Bacteria\|p__Firmicutes_D\|c__Bacilli\|o__Bacillales_G_286742\|o__Bacillales_G_286742_f__ | 5 | 11.0038168 | 0.00800424 | 0.00800424 | 5-40 |
| d__Bacteria\|p__Nitrospirota_A_437815\|c__Thermodesulfovibrionia\|o__Thermodesulfovibrionales\|o__Thermodesulfovibrionales_f__\|o__Thermode | 40 | 11.0160756 | 0.00883421 | 0.00883421 | 5-40 |
| d__Bacteria\|p__Nitrospirota_A_437815\|c__Thermodesulfovibrionia\|o__Thermodesulfovibrionales\|o__Thermodesulfovibrionales_f__ | 40 | 10.724921 | 0.00930935 | 0.00930935 | 5-40 |
| d__Bacteria\|p__Nitrospirota_A_437815\|c__Thermodesulfovibrionia\|o__Thermodesulfovibrionales\|o__Thermodesulfovibrionales_f__\|o__Thermode | 40 | 10.0392907 | 0.00992743 | 0.00992743 | 5-40 |
| d__Bacteria\|p__Actinobacteriota\|c__Acidimicrobiia_402965 | 5 | 10.7255814 | 0.00993858 | 0.00993858 | 5-40 |
| d__Bacteria\|p__Actinobacteriota\|c__Acidimicrobiia_402965\|o__UBA5794\|f__UBA5794 | 5 | 10.9195665 | 0.01086218 | 0.01086218 | 5-40 |
| d__Bacteria\|p__Actinobacteriota\|c__Acidimicrobiia_402965\|o__UBA5794 | 5 | 10.6700007 | 0.01283247 | 0.01283247 | 5-40 |
| d__Bacteria\|p__Nitrospirota_A_437815\|c__Thermodesulfovibrionia\|o__Thermodesulfovibrionales | 40 | 10.2539251 | 0.01544372 | 0.01544372 | 5-40 |
| d__Bacteria\|p__Actinobacteriota\|c__Acidimicrobiia_402965\|o__UBA5794\|f__UBA5794\|g__SZUA-442 | 5 | 11.3554943 | 0.01572668 | 0.01572668 | 5-40 |
| d__Bacteria\|p__Desulfobacterota_I\|c__Desulfobacteria\|o__Desulfobacterales | 5 | 7.67120085 | 0.02583747 | 0.02583747 | 5-40 |
| d__Bacteria\|p__Desulfobacterota_I\|c__Desulfobacteria | 5 | 7.36449881 | 0.02697684 | 0.02697684 | 5-40 |
| d__Bacteria\|p__Firmicutes_D\|c__Bacilli\|o__Bacillales_D_310495\|f__Halobacillaceae | 5 | 6.68664263 | 0.02994715 | 0.02994715 | 5-40 |
| d__Bacteria\|p__Proteobacteria\|c__Gammaproteobacteria\|o__Pseudomonadales_650611\|f__Pseudomonadaceae\|g__Pseudomonas_E_647464 | 40 | 6.93920348 | 0.03382686 | 0.03382686 | 5-40 |
| d__Bacteria\|p__Proteobacteria\|c__Gammaproteobacteria\|o__Pseudomonadales_650611\|f__Pseudomonadaceae | 40 | 6.84298573 | 0.03652031 | 0.03652031 | 5-40 |
| d__Bacteria\|p__Desulfobacterota_I\|c__Desulfobacteria\|o__Desulfobacterales\|f__SG8-13\|g__SG8-13\|s__sp001303025 | 5 | 6.07737709 | 0.04336085 | 0.04336085 | 5-40 |
| d__Bacteria\|p__Proteobacteria\|c__Gammaproteobacteria\|o__Pseudomonadales_650611\|f__Pseudomonadaceae\|g__Pseudomonas_E_647464\|s | 40 | 6.44783488 | 0.04338314 | 0.04338314 | 5-40 |
| d__Bacteria\|p__Desulfobacterota_I\|c__Desulfobacteria\|o__Desulfobacterales\|f__SG8-13 | 5 | 5.8805076 | 0.0453197 | 0.0453197 | 5-40 |
| d__Bacteria\|p__Actinobacteriota\|c__\|c___o__\|c___o___f__\|c___o___f___g__ | 5 | 5.56411815 | 0.04931592 | 0.04931592 | 5-40 |
| d__Bacteria\|p__Proteobacteria\|c__Gammaproteobacteria\|o__Pseudomonadales_641030\|f__Halomonadaceae_641030 | dry | 19.2896661 | 0.00052024 | 0.00052024 | Dry-Flood |
| d__Bacteria\|p__Proteobacteria\|c__Gammaproteobacteria\|o__Pseudomonadales_641030 | dry | 18.5417561 | 0.000551 | 0.000551 | Dry-Flood |
| d__Bacteria\|p__Proteobacteria\|c__Gammaproteobacteria\|o__Pseudomonadales_641030\|f__Halomonadaceae_641030\|g__Halomonas_E_64024 | dry | 13.6342085 | 0.00299459 | 0.00299459 | Dry-Flood |
| d__Bacteria\|p__Proteobacteria\|c__Gammaproteobacteria\|o__Pseudomonadales_641030\|f__Halomonadaceae_641030\|g__Halomonas_E_64024 | dry | 13.4044046 | 0.0052217 | 0.0052217 | Dry-Flood |
| d__Bacteria\|p__Proteobacteria\|c__Gammaproteobacteria | dry | 8.13842587 | 0.00580419 | 0.00580419 | Dry-Flood |
| d__Bacteria\|p__Proteobacteria\|c__Gammaproteobacteria\|o__Enterobacterales_A_737866 | dry | 10.5416303 | 0.0059338 | 0.0059338 | Dry-Flood |
| d__Bacteria\|p__Proteobacteria | dry | 6.30728765 | 0.01336225 | 0.01336225 | Dry-Flood |
| d__Bacteria\|p__Firmicutes_D\|c__Bacilli\|o__Lactobacillales\|f__Streptococcaceae\|g__Lactococcus_A_343306\|s__carnosus | flood | 5.61191266 | 0.02322908 | 0.02322908 | Dry-Flood |
| d__Bacteria\|p__Firmicutes_D\|c__Bacilli\|o__Lactobacillales\|f__Streptococcaceae | flood | 5.37766122 | 0.0250303 | 0.0250303 | Dry-Flood |
| d__Bacteria\|p__Proteobacteria\|c__Gammaproteobacteria\|o__Pseudomonadales_641035\|f__Marinomonadaceae | dry | 8.23120326 | 0.02552743 | 0.02552743 | Dry-Flood |
| d__Bacteria\|p__Firmicutes_D\|c__Bacilli\|o__Lactobacillales\|f__Streptococcaceae\|g__Lactococcus_A_343306 | flood | 5.2977192 | 0.02866298 | 0.02866298 | Dry-Flood |
| d__Bacteria\|p__Proteobacteria\|c__Gammaproteobacteria\|o__Pseudomonadales_641035 | dry | 7.9563284 | 0.0287004 | 0.0287004 | Dry-Flood |
| d__Bacteria\|p__Firmicutes_D\|c__Bacilli\|o__Lactobacillales | flood | 5.3688034 | 0.0318328 | 0.0318328 | Dry-Flood |
| d__Bacteria\|p__Proteobacteria\|c__Gammaproteobacteria\|o__Pseudomonadales_641035\|f__Marinomonadaceae\|g__Marinomonas\|s__piezotoler | dry | 7.59487617 | 0.03274843 | 0.03274843 | Dry-Flood |
| d__Bacteria\|p__Proteobacteria\|c__Gammaproteobacteria\|o__Enterobacterales_A_737866\|f__Vibrionaceae\|g__Vibrio_678715\|g__Vibrio_678715_ | dry | 6.43907418 | 0.03444282 | 0.03444282 | Dry-Flood |

| | | | | | |
|---|---|---|---|---|---|
| d__Bacteria\|p__Proteobacteria\|c__Gammaproteobacteria\|o__Pseudomonadales_641035\|f__Marinomonadaceae\|g__Marinomonas | dry | 7.34317539 | 0.03846903 | 0.03846903 | Dry-Flood |
| d__Bacteria\|p__Proteobacteria\|c__Gammaproteobacteria\|o__Enterobacterales_A_737866\|f__Vibrionaceae\|g__Vibrio_678715 | dry | 5.2441207 | 0.04296755 | 0.04296755 | Dry-Flood |
| d__Bacteria\|p__Firmicutes_D\|c__Bacilli\|o__Bacillales_G_286742\|o__Bacillales_G_286742_f__\|o__Bacillales_G_286742_f___g__\|o__Bacillales_G_ | dry | 5.24328879 | 0.04338366 | 0.04338366 | Dry-Flood |
| d__Bacteria\|p__Firmicutes_D\|c__Bacilli\|o__Bacillales_G_286742\|o__Bacillales_G_286742_f__ | dry | 5.46507047 | 0.04373736 | 0.04373736 | Dry-Flood |
| d__Bacteria\|p__Desulfobacterota_I\|c__Desulfobacteria\|o__Desulfobacterales | flood | 6.10583172 | 0.04535702 | 0.04535702 | Dry-Flood |
| d__Bacteria\|p__Proteobacteria\|c__Gammaproteobacteria\|o__Enterobacterales_A_737866\|f__Vibrionaceae | dry | 5.23033002 | 0.04544262 | 0.04544262 | Dry-Flood |
| d__Bacteria\|p__Firmicutes_D\|c__Bacilli\|o__Bacillales_G_286742\|o__Bacillales_G_286742_f__\|o__Bacillales_G_286742_f___g__ | dry | 5.45649044 | 0.04583426 | 0.04583426 | Dry-Flood |
| d__Bacteria\|p__Proteobacteria\|c__Gammaproteobacteria\|o__Enterobacterales_A_737866\|f__Alteromonadaceae_665222 | dry | 6.64379268 | 0.04602466 | 0.04602466 | Dry-Flood |
| d__Bacteria\|p__Firmicutes_D\|c__Bacilli\|o__Bacillales_G_286742 | dry | 5.06513376 | 0.04726614 | 0.04726614 | Dry-Flood |
| d__Bacteria\|p__Chloroflexota\|c__Anaerolineae\|o__Anaerolineales | Impaired | 26.1911739 | 0.0002223 | 0.0002223 | MD-D |
| d__Bacteria\|p__Chloroflexota\|c__Anaerolineae\|o__Anaerolineales\|f__UBA4823 | Impaired | 20.9030673 | 0.00081603 | 0.00081603 | MD-D |
| d__Bacteria\|p__Firmicutes_D\|c__Bacilli\|o__Lactobacillales\|f__Streptococcaceae\|g__Lactococcus_A_343306\|s__carnosus | Basin | 12.1817215 | 0.00101782 | 0.00101782 | MD-D |
| d__Bacteria\|p__Firmicutes_D\|c__Bacilli\|o__Lactobacillales | Basin | 11.983995 | 0.00108674 | 0.00108674 | MD-D |
| d__Bacteria\|p__Firmicutes_D\|c__Bacilli\|o__Lactobacillales\|f__Streptococcaceae\|g__Lactococcus_A_343306 | Basin | 11.9590482 | 0.00110129 | 0.00110129 | MD-D |
| d__Bacteria\|p__Firmicutes_D\|c__Bacilli\|o__Lactobacillales\|f__Streptococcaceae | Basin | 11.842291 | 0.00111798 | 0.00111798 | MD-D |
| d__Bacteria\|p__Chloroflexota\|c__Anaerolineae | Impaired | 14.5817732 | 0.00137584 | 0.00137584 | MD-D |
| d__Bacteria\|p__Nitrospirota_A_437815 | Impaired | 8.89567587 | 0.01087259 | 0.01087259 | MD-D |
| d__Bacteria\|p__Chloroflexota\|c__Anaerolineae\|o__Anaerolineales\|f__UBA4823\|g__DSWF01\|s__sp011368195 | Impaired | 9.67062675 | 0.01232308 | 0.01232308 | MD-D |
| d__Bacteria\|p__Nitrospirota_A_437815\|c__Thermodesulfovibrionia\|c__Thermodesulfovibrionia_o__\|c__Thermodesulfovibrionia_o___f__\|c__Therm | Impaired | 8.78073704 | 0.01443375 | 0.01443375 | MD-D |
| d__Bacteria\|p__Nitrospirota_A_437815\|c__Thermodesulfovibrionia\|c__Thermodesulfovibrionia_o__\|c__Thermodesulfovibrionia_o___f__ | Impaired | 9.34492442 | 0.01452266 | 0.01452266 | MD-D |
| d__Bacteria\|p__Nitrospirota_A_437815\|c__Thermodesulfovibrionia | Impaired | 8.44955897 | 0.01606576 | 0.01606576 | MD-D |
| d__Bacteria\|p__Chloroflexota\|c__Anaerolineae\|o__Anaerolineales\|f__UBA4823\|g__DSWF01 | Impaired | 10.2619882 | 0.01642812 | 0.01642812 | MD-D |
| d__Bacteria\|p__Proteobacteria\|c__Gammaproteobacteria\|o__Pseudomonadales_641030\|f__Halomonadaceae_641030 | Basin | 7.64628785 | 0.0188222 | 0.0188222 | MD-D |
| d__Bacteria\|p__Nitrospirota_A_437815\|c__Thermodesulfovibrionia\|c__Thermodesulfovibrionia_o__ | Impaired | 8.55400414 | 0.02043346 | 0.02043346 | MD-D |
| d__Bacteria\|p__Proteobacteria\|c__Gammaproteobacteria\|o__Pseudomonadales_641030 | Basin | 7.76288316 | 0.02074967 | 0.02074967 | MD-D |
| d__Bacteria\|p__Firmicutes_D | Basin | 5.66660615 | 0.02155848 | 0.02155848 | MD-D |
| d__Bacteria\|p__Firmicutes_D\|c__Bacilli | Basin | 5.49905405 | 0.02222861 | 0.02222861 | MD-D |
| d__Bacteria\|p__Nitrospirota_A_437815\|c__Thermodesulfovibrionia\|c__Thermodesulfovibrionia_o__\|c__Thermodesulfovibrionia_o___f__\|c__Therm | Impaired | 8.64775145 | 0.02321141 | 0.02321141 | MD-D |
| d__Bacteria\|p__Firmicutes_D\|c__Bacilli\|o__Bacillales_B_302584\|f__Bacillaceae_B_301660\|g__Rossellomorea\|g__Rossellomorea_s__ | Basin | 7.97078252 | 0.02575403 | 0.02575403 | MD-D |
| d__Bacteria\|p__Firmicutes_D\|c__Bacilli\|o__Bacillales_B_302584\|f__Bacillaceae_B_301660\|g__Rossellomorea | Basin | 8.02867812 | 0.02887659 | 0.02887659 | MD-D |
| d__Bacteria\|p__Proteobacteria\|c__Gammaproteobacteria\|o__Pseudomonadales_641030\|f__Halomonadaceae_641030\|g__Halomonas_E_64024 | Basin | 5.86608798 | 0.04588829 | 0.04588829 | MD-D |
| d__Bacteria\|p__Proteobacteria\|c__Gammaproteobacteria\|o__Pseudomonadales_641030\|f__Halomonadaceae_641030\|g__Halomonas_E_64024 | Basin | 5.60454898 | 0.04612508 | 0.04612508 | MD-D |
| d__Bacteria\|p__Chloroflexota | Impaired | 29.5513571 | 1.34E-05 | 1.34E-05 | MD-D |
| d__Bacteria\|p__Desulfobacterota_I\|c__DSM-4660\|o__Desulfatiglandales | Fringe | 7.74865874 | 0.02424244 | 0.02424244 | MD-ND |
| d__Bacteria\|p__Chloroflexota\|c__Anaerolineae\|o__Anaerolineales | Fringe | 7.83512683 | 0.02719146 | 0.02719146 | MD-ND |
| d__Bacteria\|p__Desulfobacterota_I\|c__DSM-4660\|o__Desulfatiglandales\|f__HGW-15\|g__JAAZBV01\|g__JAAZBV01_s__ | Fringe | 7.35673595 | 0.02836686 | 0.02836686 | MD-ND |
| d__Bacteria\|p__Desulfobacterota_I\|c__DSM-4660 | Fringe | 7.02148596 | 0.03129516 | 0.03129516 | MD-ND |
| d__Bacteria\|p__Desulfobacterota_I\|c__DSM-4660\|o__Desulfatiglandales\|f__HGW-15 | Fringe | 7.0627874 | 0.03435065 | 0.03435065 | MD-ND |
| d__Bacteria\|p__Desulfobacterota_I\|c__DSM-4660\|o__Desulfatiglandales\|f__HGW-15\|g__JAAZBV01 | Fringe | 6.58200625 | 0.03470354 | 0.03470354 | MD-ND |
| d__Bacteria\|p__Proteobacteria\|c__Gammaproteobacteria\|o__Enterobacterales_A_737866\|f__Vibrionaceae\|g__Vibrio_678715\|s__Vibrio rumoiens | Fringe | 5.47685344 | 0.04778032 | 0.04778032 | MD-ND |
| d__Bacteria\|p__Chloroflexota | Impaired | 11.944607 | 0.00242985 | 0.00242985 | ND-D |
| d__Bacteria\|p__Proteobacteria\|c__Alphaproteobacteria | Impaired | 11.0481877 | 0.00544347 | 0.00544347 | ND-D |
| d__Bacteria\|p__Desulfobacterota_I\|c__DSM-4660\|o__Desulfatiglandales\|f__HGW-15\|g__JAAZBV01 | Fringe | 11.7896006 | 0.00758184 | 0.00758184 | ND-D |
| d__Bacteria\|p__Desulfobacterota_I\|c__DSM-4660\|o__Desulfatiglandales\|f__HGW-15\|g__JAAZBV01\|g__JAAZBV01_s__ | Fringe | 11.2741181 | 0.00860408 | 0.00860408 | ND-D |
| d__Bacteria\|p__Firmicutes_D\|c__Bacilli\|o__Bacillales_B_302584\|f__Bacillaceae_B_301660\|g__Rossellomorea\|g__Rossellomorea_s__ | Fringe | 9.33020563 | 0.01666848 | 0.01666848 | ND-D |
| d__Bacteria\|p__Firmicutes_D\|c__Bacilli\|o__Bacillales_B_302584\|f__Bacillaceae_B_301660\|g__Rossellomorea | Fringe | 9.78270238 | 0.01678819 | 0.01678819 | ND-D |
| d__Bacteria\|p__Proteobacteria\|c__Gammaproteobacteria\|o__Enterobacterales_A_737866\|f__Vibrionaceae\|g__Vibrio_678715\|s__Vibrio rumoiens | Fringe | 8.14200689 | 0.01781567 | 0.01781567 | ND-D |
| d__Bacteria\|p__Chloroflexota\|c__Anaerolineae\|o__Anaerolineales\|f__UBA4823\|g__DSWF01 | Impaired | 9.16135989 | 0.0191867 | 0.0191867 | ND-D |
| d__Bacteria\|p__Proteobacteria\|c__Gammaproteobacteria\|o__Enterobacterales_A_737866\|f__Vibrionaceae | Fringe | 6.52363768 | 0.02327382 | 0.02327382 | ND-D |

<table>
<tr><td>d__Bacteria|p__Chloroflexota|c__Anaerolineae|o__Anaerolineales|f__UBA4823|g__DSWF01|s__sp011368195</td><td>Impaired</td><td>8.26201591</td><td>0.02465924</td><td>0.02465924</td><td>ND-D</td></tr>
<tr><td>d__Bacteria|p__Nitrospirota_A_437815|c__Thermodesulfovibrionia|c__Thermodesulfovibrionia_o__|c__Thermodesulfovibrionia_o___f__</td><td>Impaired</td><td>7.59384236</td><td>0.02853199</td><td>0.02853199</td><td>ND-D</td></tr>
<tr><td>d__Bacteria|p__Proteobacteria|c__Gammaproteobacteria|o__Enterobacterales_A_737866|f__Vibrionaceae|g__Vibrio_678715|g__Vibrio_678715_</td><td>Fringe</td><td>7.95575287</td><td>0.03035386</td><td>0.03035386</td><td>ND-D</td></tr>
<tr><td>d__Bacteria|p__Chloroflexota|c__Anaerolineae</td><td>Impaired</td><td>5.67328538</td><td>0.03076792</td><td>0.03076792</td><td>ND-D</td></tr>
<tr><td>d__Bacteria|p__Nitrospirota_A_437815|c__Thermodesulfovibrionia</td><td>Impaired</td><td>6.16390003</td><td>0.03132088</td><td>0.03132088</td><td>ND-D</td></tr>
<tr><td>d__Bacteria|p__Nitrospirota_A_437815</td><td>Impaired</td><td>5.89664169</td><td>0.03134598</td><td>0.03134598</td><td>ND-D</td></tr>
<tr><td>d__Bacteria|p__Nitrospirota_A_437815|c__Thermodesulfovibrionia|c__Thermodesulfovibrionia_o__|c__Thermodesulfovibrionia_o___f__|c__Therm</td><td>Impaired</td><td>7.56536217</td><td>0.03385473</td><td>0.03385473</td><td>ND-D</td></tr>
<tr><td>d__Bacteria|p__Nitrospirota_A_437815|c__Thermodesulfovibrionia|c__Thermodesulfovibrionia_o__</td><td>Impaired</td><td>7.42575345</td><td>0.03456936</td><td>0.03456936</td><td>ND-D</td></tr>
<tr><td>d__Bacteria|p__Proteobacteria|c__Gammaproteobacteria|o__Enterobacterales_A_737866|f__Vibrionaceae|g__Vibrio_678715</td><td>Fringe</td><td>6.19891331</td><td>0.0382127</td><td>0.0382127</td><td>ND-D</td></tr>
<tr><td>d__Bacteria|p__Nitrospirota_A_437815|c__Thermodesulfovibrionia|c__Thermodesulfovibrionia_o__|c__Thermodesulfovibrionia_o___f__|c__Therm</td><td>Impaired</td><td>6.39505944</td><td>0.03874158</td><td>0.03874158</td><td>ND-D</td></tr>
<tr><td>d__Bacteria|p__Desulfobacterota_I|c__DSM-4660|o__Desulfatiglandales|f__HGW-15</td><td>Fringe</td><td>6.48639027</td><td>0.03976244</td><td>0.03976244</td><td>ND-D</td></tr>
<tr><td>d__Bacteria|p__Bacteroidota|c__Bacteroidia|o__Bacteroidales</td><td>Impaired</td><td>6.21870791</td><td>0.0434178</td><td>0.0434178</td><td>ND-D</td></tr>
<tr><td>d__Bacteria|p__Firmicutes_D</td><td>Fringe</td><td>4.12681821</td><td>0.04458125</td><td>0.04458125</td><td>ND-D</td></tr>
<tr><td>d__Bacteria|p__Firmicutes_D|c__Bacilli</td><td>Fringe</td><td>4.09153566</td><td>0.04565264</td><td>0.04565264</td><td>ND-D</td></tr>
</table>
