## Supplementary material for "Flooding patterns shape microbial community in mangrove sediments": Supplemantary Table 2

**Supplementary Table 2. PERMANOVA Analysis of PCoA Using Bray-Curtis Distance.**

Permutation test for adonis under reduced model
Marginal effects of terms
Permutation: free
Number of permutations: 9999

vegan::adonis2(formula = formula, data = metadata, permutations = n_perms, by = by, parallel = parall)

| Term | Df | Sum of Squa | $R^2$ | F | Pr(>F) | Significance |
|---|---|---|---|---|---|---|
| Zone | 2 | 1.7244 | 0.07264 | 2.0803 | 1.00E-04 | *** |
| Season | 1 | 0.8382 | 0.03531 | 2.0223 | 2.00E-04 | *** |
| Depth | 2 | 1.7089 | 0.07199 | 2.0616 | 1.00E-04 | *** |
| Residual | 47 | 19.4792 | 0.8206 | | | |
| Total | 52 | 23.7378 | 1 | | | |

**Significance codes:**
*** = 0.001
** = 0.01

0.05
. = 0.1
' ' = 1
