## Supplementary Table 3 for "Flooding patterns shape microbial community in mangrove sediments"

**Supplementary Table 3. PERMANOVA Analysis of PCoA Using UniFrac Distance.**

Permutation test for adonis under reduced model
Marginal effects of terms
Permutation: free
Number of permutations: 9999

vegan::adonis2(formula = formula, data = metadata, permutations = n_perms, by = by, parallel = parall)

| Term | Df | Sum of Squar | $R^2$ | F | Pr(>F) | Significance |
|---|---|---|---|---|---|---|
| Zone | 2 | 0.9636 | 0.13314 | 4.3877 | 1.00E-04 | *** |
| Season | 1 | 0.4574 | 0.0632 | 4.1658 | 1.00E-04 | *** |
| Depth | 2 | 0.6598 | 0.09117 | 3.0045 | 1.00E-04 | *** |
| Residual | 47 | 5.1611 | 0.71308 | | | |
| Total | 52 | 7.2377 | 1 | | | |

**Significance codes:**
*** = 0.001
** = 0.01

0.05
. = 0.1
' ' = 1
